# Laboratory evaluation of a quaternary ammonium compound (QAC)-based antimicrobial coating used in public transport during the COVID-19 pandemic

**DOI:** 10.1101/2022.10.12.512011

**Authors:** Paz Aranega-Bou, Natalie Brown, Abigail Stigling, Wilhemina D’Costa, Neville Q Verlander, Thomas Pottage, Allan Bennett, Ginny Moore

## Abstract

The virucidal activity of a quaternary ammonium compound (QAC)-based antimicrobial coating used by the UK rail industry during the COVID-19 pandemic was evaluated using the bacteriophage ϕ6 as a surrogate for SARS-CoV-2. Immediately after application and in the absence of interfering substance, the product showed efficacy (>3 log_10_ reduction) on some materials typically used in rail carriages (stainless steel, high pressure laminate and plastic), variable efficacy on glass and no efficacy (<3 log_10_ reduction) on a train armrest made of Terluran 22. If, after application of the product, the surfaces remained undisturbed, the antimicrobial coating retained its efficacy for at least 28 days on all materials where it was effective immediately after application. However, regardless of the material coated or time since application, the presence of organic debris (fetal bovine serum) significantly reduced the viricidal activity of the coating. Wiping the surface with a wetted cloth after organic debris deposition was not sufficient to restore efficacy. We conclude that the product is likely to be of limited effectiveness in a busy multi-user environment such as public transport.

**Importance:** This study evaluated the performance of a commercially available antimicrobial coating used by the transport industry in the UK during the COVID-19 pandemic. While the product initially showed efficacy against ϕ6 when applied to some materials, when organic debris was subsequently deposited, the efficacy was severely diminished and could not be recovered through wiping (cleaning) the surface. This highlights the importance of including relevant materials and conditions when evaluating antimicrobial coatings in the laboratory. Further efforts are required to identify suitable infection prevention and control practices for the transport industry.

## Introduction

Surfaces can become contaminated with virus via direct contact with body fluids or soiled hands and via deposition of aerosolized virus. Contaminated surfaces can act as fomites and contribute to the spread of viral infections (1) and, while fomite transmission is not currently believed to be the primary transmission route for SARS-CoV-2 (2), the importance of this pathway in relation to other pathways might differ in different venues and situations (3). While the risk of transmission following brief contact with a single contaminated surface is estimated to be low, the risk increases as individuals touch a higher number of contaminated surfaces (2). Factors such as the number of contaminated hand contact sites, the density of individuals and duration of stay within a venue, along with any mitigation strategies, such as surface decontamination and handwashing, influence the risk of transmission of viruses via fomites (2, 3). Use of public transport can require an individual to touch or grip a number of surfaces, including poles, seat head rests, tables, push buttons and armrests.

Despite usage remaining considerably lower than pre-pandemic levels, during the 2020-2021 financial year, 388 million rail passenger journeys were made in Great Britain increasing to 990 million during 2021-22 (4) when 1.57 billion local bus passenger journeys were made in England alone (5). Public transport use and accessibility is associated with better air quality, higher rates of employment and lower social exclusion and can encourage a more active lifestyle (6). However, the role that public transport plays in transmission of infectious diseases is not well understood although there is some epidemiological evidence that it could contribute to transmission of influenza-like illness (7, 8) and high-touch surfaces on public transport vehicles are known to be contaminated with bacteria (9) and can be contaminated with SARS-CoV-2 RNA (10, 11). A recent modelling study comparing the relative contributions of close-range exposure to SARS-CoV-2 (via droplets or aerosols), airborne exposure (via small aerosols without having to be within 2 metres of the infectious source) and exposure via contaminated fomites in a subway carriage concluded that all three routes of transmission are relevant in this setting (12).

The field of antimicrobial coatings has developed rapidly in recent years and their use (in conjunction with handwashing and cleaning) as a potential infection prevention strategy to reduce healthcare-associated infections has been described (13). The COVID-19 pandemic has demonstrated the need to apply infection prevention and control principles in public spaces outside healthcare, such as public transport. However, guidelines are scarce, and more research is needed to understand what practices are appropriate for the transport sector.

In the UK, some transport operators, in addition to implementing enhanced cleaning protocols, have used antimicrobial coatings in an attempt to reduce surface bioburden and the risk of viral contamination. There are a number of products commercially available that can be applied to existing surfaces and, according to manufacturers’ claims, provide long lasting residual antimicrobial activity when they are used in conjunction with regular cleaning (14). The US Environmental Protection Agency (US EPA) has developed an interim method that outlines the requirements for the registration of antimicrobial coatings that are intended to provide residual antimicrobial activity for a period of weeks and are applied to surfaces to supplement standard disinfection practices. The protocol includes efficacy assessment after coated surfaces are subjected to cycles of dry and wet abrasion using specialised equipment to simulate wear (15). However, this equipment is not widely available and, in most cases, manufacturer’s claims are supported by limited evidence, often obtained through laboratory tests that have not attempted to mimic real-life conditions. Here we used the bacteriophage ϕ6 as a surrogate for SARS-CoV-2 to evaluate the antiviral efficacy of Zoono Z71 Microbe Shield Surface Sanitiser and Protectant (Zoono™, Bury St Edmunds, UK) which has been used by the transport industry throughout the pandemic.

## Methods

### Virus propagation

Bacteriophage ϕ6 (DSM 21518) and host bacteria *Pseudomonas syringae* (*Pseudomonas* sp. (DSM 21482)) were obtained from the DSMZ GmbH where they had been deposited by S. Moineau from Université Laval, Québec. *P. syringae* was reconstituted according to the manufacturer instructions, stored on cryobeads (Technical Service Consultants, Heywood, UK) at -80°C and cultured weekly on Tryptone Soya Agar (TSA) (Oxoid Ltd, Basingstoke, UK). ϕ6 was stored at 4 °C (for no longer than 5 months) and propagated in *P. syringae* host cells.

A 10 µL loopful of *P. syringae* was transferred from TSA to 10 mL Tryptone Soya Broth (TSB; E&O Laboratories Ltd, Bonnybridge, UK) and incubated for 18-20 hours (25 °C; 150 rpm). 100 µL of the resulting suspension were transferred to 10 mL TSB and incubated at 25 °C (170 rpm). When the turbidity of the broth culture (OD_600_) exceeded 0.1 (approx. 5 hours), 50 µL of ϕ6 suspension was added. Cultures were incubated overnight (25 °C; 150-170 rpm) until total lysis. Propagated phage was filtered using a 0.22 µm PES syringe filter (Starlab, Milton Keynes, UK) and the concentration was determined via plaque assay. Phage suspensions (∼2×10^10^ pfu/mL) were stored at 4°C for up to 5 months.

### Preparation and coating of test coupons

Glass coupons (1.56 cm^2^) were cut from microscope slides (Marienfeld, Lauda-Königshofen, Germany) using a glass cutter. Polystyrene coupons (2.54 cm^2^) were bought from Amazon (Fingooo or Suloli brands). Stainless steel coupons (1.44 cm^2^) were obtained from Apsley Precision Engineering Ltd (Salisbury, UK). Glass and stainless-steel coupons were washed with 5% Decon 90 (Decon, Sussex, UK) for 5 minutes, rinsed with demineralised water and 70% isopropanol (IPA) allowed to dry and then autoclaved (121 °C for 15 minutes). Polystyrene coupons were disinfected with 70% IPA and left to dry. The product information sheet lists the quaternary ammonium compound Alkyl (C12-16) dimethylbenzyl ammonium chloride (ADBAC/BKC (C12-16)) CAS 68424-85-1 (commonly known as Benzalkonium chloride) at a concentration of 0.1% (w/w) as the active ingredient of Zoono Z71. Glass coupons were initially coated by manual spraying but, due to low reproducibility, it was decided to pipette 75 µL of the product on to each coupon and to spread it evenly over the surface. Stainless steel and polystyrene coupons were coated by pipetting 40 µL per coupon. The product was applied ∼24 hours before inoculation unless otherwise stated.

### Preparation and coating of train parts

Previously used train parts were donated by First Group (Aberdeen, UK) from a scrap train. These consisted of tray tables made of high pressure laminate and stainless steel, armrests made of Terluran 22, and hand poles with a plastic coating (Figure 1). Upon receipt, train parts were cleaned and disinfected by wiping with 70% IPA wipes (Sani-Cloth) until wipes were not visibly dirty after the surface was wiped. Between experiments, train parts were cleaned with 70% IPA and a Kimtech™ 7644 wiper, before a further neutralisation step using Dey-Engley neutralizing broth (DEB) (Sigma-Aldrich/Merck) and then rinsed with sterile water. The last two steps were to ensure the antimicrobial coating previously applied had been neutralised and removed. The coating was applied using an electrostatic sprayer (Comac E-Spray) to ensure a homogenous distribution. Train parts were placed in a Class III cabinet, with the front panel removed and fans on whilst spraying was undertaken. Train parts were sprayed from a distance of ∼0.15-0.3m until the surfaces were thoroughly wetted, and then allowed to dry overnight within the cabinet without the fan operating.

**Figure 1:**
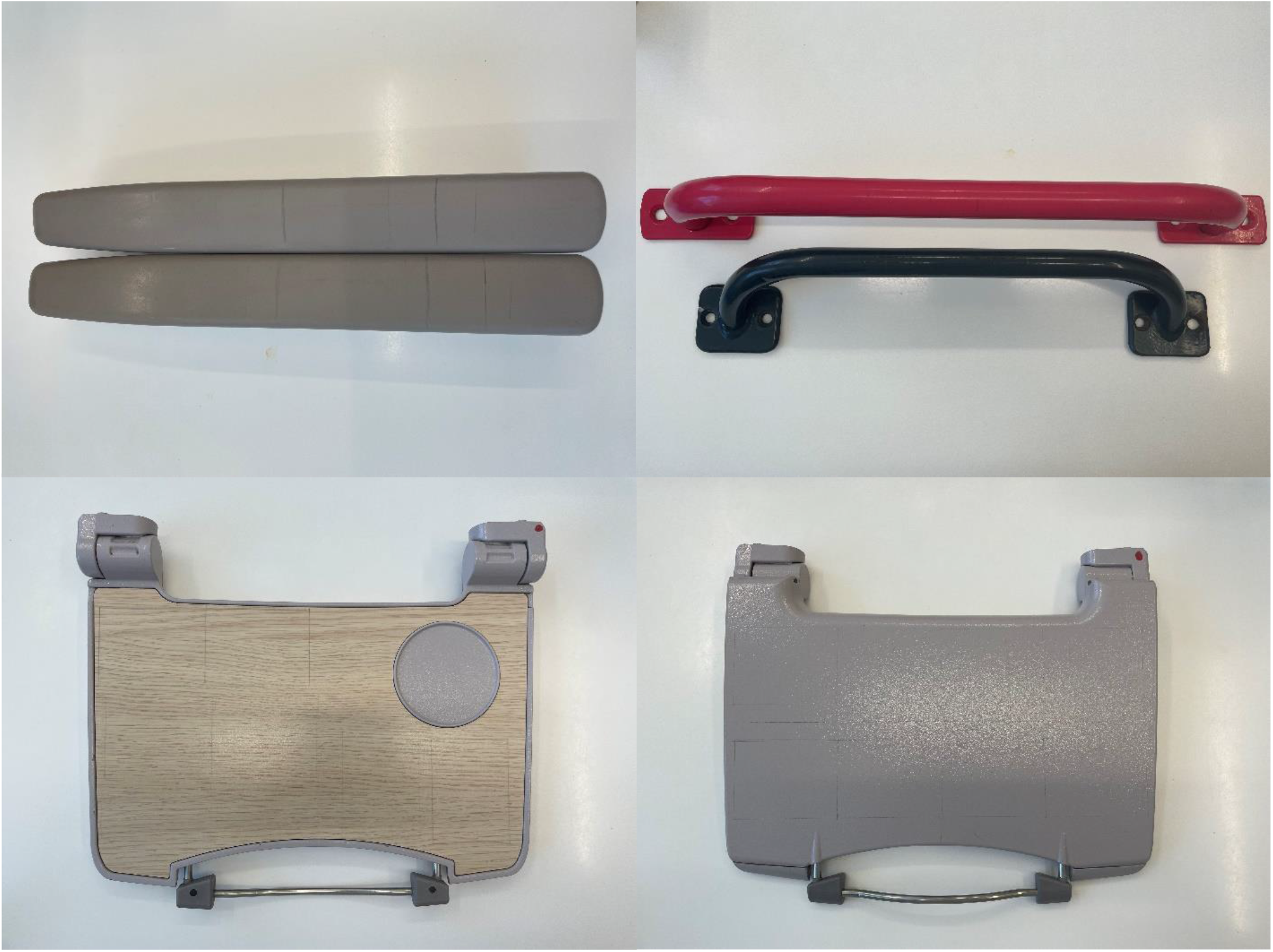
Top left – Terluran 22 armrests; top right – plastic coated hand poles; bottom left – high pressure laminate tray table; bottom right – stainless steel tray table.

### Efficacy testing

Coated and non-coated test surfaces (material coupons and train parts) were inoculated with 10 µL droplets of a ϕ6 suspension (∼2 × 10^8^ pfu) with the exception of the hand pole which was inoculated with 5 µL droplets due to structural constraints (∼1 × 10^8^ pfu). Surfaces were sampled either immediately (t=0) or after a pre-determined contact time of up to 120 minutes under ambient laboratory conditions. At each time point, individual coupons (n=3) were transferred to tubes containing 2mL DEB and four glass beads. Train parts (inoculated with multiple droplets) were sampled using FLOQSwabs™ (Copan Diagnostics, Murrieta, USA). Three test areas were swabbed at each time point and each swab was placed in 2mL DEB containing four glass beads. The suitability of DEB to neutralise the antimicrobial coating in liquid form was assessed before any efficacy tests were carried out (16). Tubes were vortexed for 30 seconds and serially diluted in TSB before the concentration of ϕ6 was determined via plaque assay.

### Plaque assay

Soft phage agar (0.6%; Media Services, UKHSA) in 3 mL aliquots was melted (by heating to 100 °C), maintained at 60 °C and cooled (37 °C, 5 min) before being mixed with 250 µL *P syringae* (grown in TSB; OD_600_ > 0.9) and 100 µL of test suspension. The soft phage agar was then poured over the surface of a TSA plate and evenly distributed. Once the soft agar layer had set, the plates were incubated at 25 °C for 18 – 24 h and the resulting plaques manually enumerated. Assays were conducted in duplicate and included control samples containing no ϕ6 to monitor for contamination and to ensure appropriate lawn formation by *P. syringae*. Validation work showed that samples could be kept at room temperature for three hours and at 4 °C for up to a week with minimal changes in concentration (average log_10_ change from -0.07 to 0.05) (Supplementary table 2). Based on these results, samples were stored at 4 °C for up to a week after collection and plaque assays repeated when *P. syringae* lawns did not show strong growth. Efficacy was calculated by subtracting the average log_10_ concentration at any given timepoint from the average log_10_ concentration immediately after inoculation (t=0). The coating was considered effective against ϕ6 if, under any given set of conditions, the log_10_ reduction achieved was ≥ 3.0. This criterion was based on the US EPA interim guidance for evaluating the efficacy of antimicrobial surface coatings, which states that a 3 log_10_ reduction of test microbes within 1–2 hours contact times is required for product registration (15). Non-coated coupons/train parts were always processed in parallel to monitor survival of ϕ6 over the course of the experiment. Surfaces were coated up to 28 days prior to inoculation to assess claims that efficacy is retained for 30 days post-application.

### Application of interfering substances

To mimic contamination with organic matter, 40µL of bovine serum albumin (BSA 3 g/L; Sigma-Aldrich) or fetal bovine serum (FBS; Sigma-Aldrich) was applied to coated and non-coated coupons prior to inoculation with ϕ6. Efficacy was then assessed on coated and non-coated coupons (with and without interfering substances) as described in the previous section.

Similarly, areas of 48, 45 and 25.5 cm^2^ were drawn out on coated and non-coated tray tables, arm rests and poles respectively. FBS was applied to each test area (200 µL for tray tables and arm rests, and 100 µL for hand poles) and spread across the whole area using FLOQSwabs™ (Copan Diagnostics). The train parts were left to dry under ambient laboratory conditions before inoculation with ϕ6. The survival of ϕ6 on coated and non-coated test areas (with and without FBS) was assessed.

### Wiping

A wiping protocol was developed to simulate the mechanical action of cleaning. J cloths (Chicopee® J-Cloth Plus) were cut into 48 cm^2^ swatches and moistened with sterile water. They were then used to wipe the surface of a tray table upwards 5 times and across 5 times (10 wipe protocol) or upwards 20 times and across 20 times (40 wipe protocol; only applied to the high pressure laminate side). Both coated and non-coated tray tables (in the presence and absence of dried FBS) were subjected to wiping. Following wiping, the survival of ϕ6 on coated and non-coated test surfaces was assessed as described previously. Experiments were carried out in duplicate with three technical replicates per experiment.

### Statistical analysis

The statistical analysis was done with a regression framework wherein statistical significance was set at 5% and the composite Wald test used to ascertain statistical significance. Details about the models, the estimates and their 95% Confidence Intervals (CIs) and p-values are available in the supplementary material. All the analysis was carried out in STATA 17.0.

## Results

### Efficacy testing on coupons

Under laboratory conditions and in the absence of interfering substance, Zoono Z71 was effective against ϕ6 (i.e. achieved ≥3 log_10_ reduction of viability) when applied to stainless steel and polystyrene. When applied to glass, the coating reduced the mean concentration of ϕ6 by 1.4 and 4.3 log_10_ values over the 30- and 60-minute contact times respectively (n=9). However, this reduction was variable and ranged from -0.1 to 6.0 and from 2.9 to 6.0 log_10_ values for individual coupons sampled after 30- and 60-minutes respectively (Figure 2). On non-coated glass coupons, losses in ϕ6 viability were minimal (0.0 to 0.2 log_10_ values), indicating that ϕ6 was stable on the coupons over the course of the experiment and natural losses in viability were not the reason for the variability observed (Figure 2). The reduction of ϕ6 concentration on coated glass surfaces was statistically significant but the two methods of application (manual spraying and pipetting) did not significantly differ from each other (Supplementary table 3).

**Figure 2:**
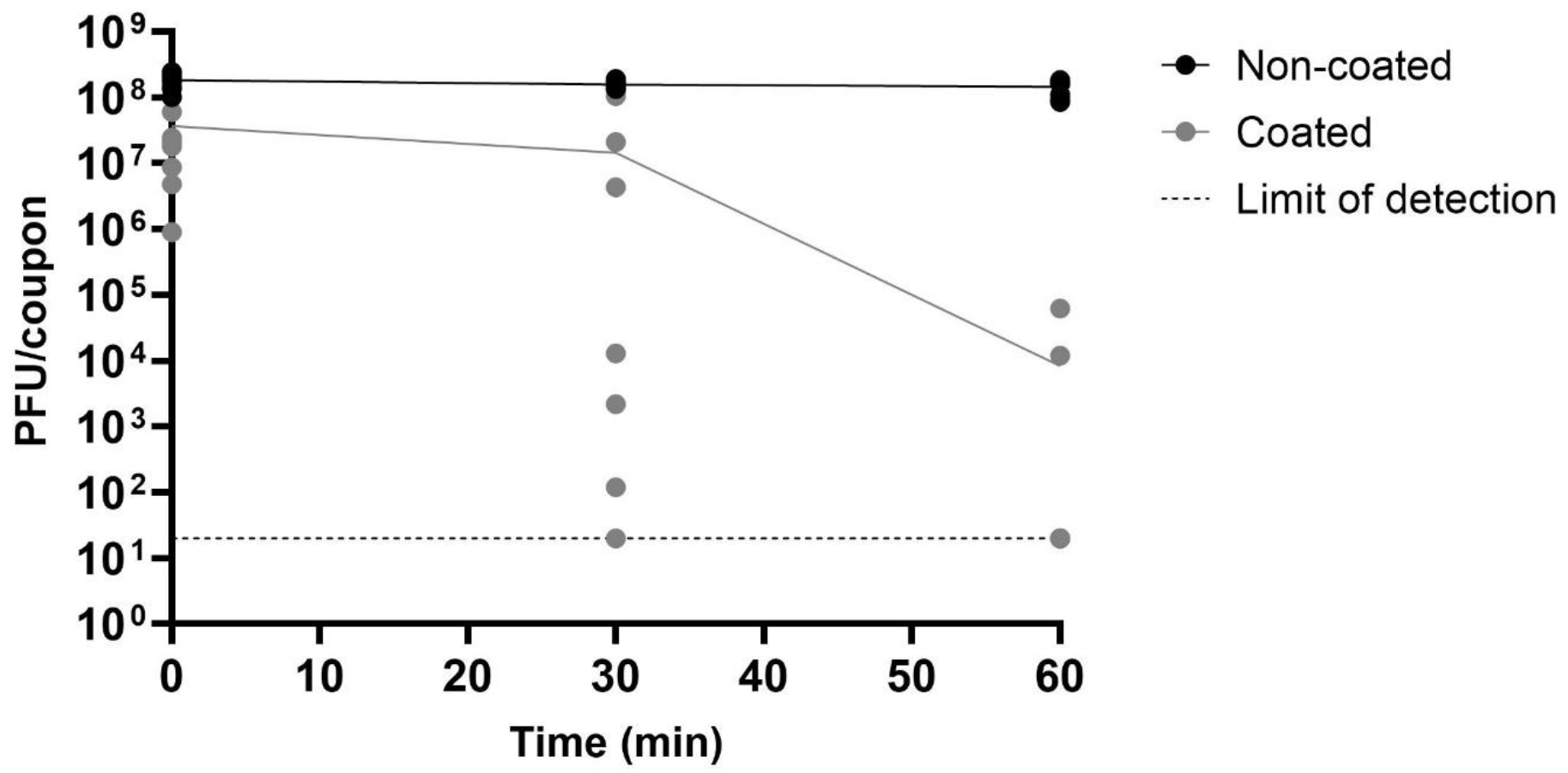
Survival of ɸ6 over 60 minutes on glass coupons coated (grey circles) with Zoono Z71 and non-coated (black circles) coupons processed in parallel. Coupons were coated by either spraying (n=6) or pipetting 75 µL on each coupon (n=3). All conditions were tested in triplicate on three separate occasions and individual results expressed as plaque forming units per coupon (PFU/coupon). Individual dots represent the results for individual coupons and lines show the average for each time point. No ϕ6 could be recovered from three out of nine coated coupons after 30 minutes and seven out of nine coated coupons after 60 minutes. The concentration of these samples was assumed to be 20 PFU/coupon, the theoretical limit of detection, for analysis.

In contrast, Zoono Z71 achieved >6.0 log_10_ reductions on all coupons within a 15-minute contact time when applied to stainless steel or polystyrene, irrespective of number of days since application, with the exception of stainless steel at day 28 post application where 30 minutes of contact time was required (Table 1). Under the experimental conditions described (no disturbance of the coupons or interfering material present), neither material type (stainless steel or polystyrene) nor number of days since application of the coating had a significant effect on efficacy (Supplementary table 4).

**Table 1:** Efficacy of Zoono Z71 when applied to stainless steel or polystyrene surfaces and inoculated with ɸ6. Efficacy is expressed as mean log10 reduction (n=3) calculated by subtracting the mean ɸ6 pfu recovered at each contact time from mean ɸ6 pfu recovered immediately after inoculation (time=0). Undetected virus observations were assumed to have a concentration of 20 PFU/coupon, the theoretical limit of detection, for analysis.

| Days after coating was applied | Stainless steel |  |  |  |  | Polystyrene |  |  |  |  |
| --- | --- | --- | --- | --- | --- | --- | --- | --- | --- | --- |
|  | Contact time |  |  |  |  |  |  |  |  |  |
|  | 5 min | 15 min | 30 min | 60 min | 120 min | 5 min | 15 min | 30 min | 60 min | 120 min |
| 1 | 0.8 | ≥6.2 | ≥6.2 | ≥6.2 | ≥6.2 | 5.9 | ≥6.0 | ≥6.0 | ≥6.0 | ≥6.0 |
| 7 | 1.4 | ≥6.8 | ≥6.8 | ≥6.8 | ≥6.8 | 3.4 | ≥6.5 | ≥6.5 | ≥6.5 | ≥6.5 |
| 14 | 1.8 | ≥6.6 | ≥6.6 | ≥6.6 | ≥6.6 | 4.0 | ≥6.5 | ≥6.5 | ≥6.5 | ≥6.5 |
| 21 | 5.9 | ≥6.5 | ≥6.5 | ≥6.5 | ≥6.5 | 0.4 | ≥6.3 | ≥6.3 | ≥6.3 | ≥6.3 |
| 28 | -0.1 | 0.3 | ≥6.7 | ≥6.7 | ≥6.7 | 3.6 | ≥6.6 | ≥6.6 | ≥6.6 | ≥6.6 |

### Evaluation of BSA and FBS as interfering substances

To mimic the accumulation of organic material on surfaces present in busy spaces such as public transport, a layer of BSA or FBS was applied to coated and non-coated coupons prior to the inoculation of ϕ6. When FBS was applied over the antimicrobial coating the average log_10_ counts were significantly higher compared to when BSA or no interfering material was applied, with no difference found between the latter two conditions (Supplementary table 5). In the presence of FBS, contact times of at least 60 minutes were required to achieve >3 log_10_ reductions while 15 minutes was required in the presence of BSA (Table 2).

**Table 2:** Efficacy of Zoono ZT1 when applied to stainless steel and polystyrene test surfaces. Surfaces were coated with a layer of BSA or FBS prior to inoculation of ϕ6 and efficacy (calculated as mean log1_0_ reduction (n=3)) was compared to when no interfering substance (clean) was present. Undetected virus observations were assumed to have a concentration of 20 PFU/coupon, the theoretical detection limit, for analysis.

|  | Stainless steel |  |  |  | Polystyrene |  |  |  |
| --- | --- | --- | --- | --- | --- | --- | --- | --- |
|  | Contact time |  |  |  |  |  |  |  |
|  | 15 min | 30 min | 60 min | 120 min | 15 min | 30 min | 60 min | 120 min |
| Clean | 4.3 | ≥6.2 | ≥6.2 | ≥6.2 | ≥6.8 | ≥6.8 | ≥6.8 | ≥6.8 |
| BSA | 6.2 | ≥6.5 | ≥6.5 | ≥6.5 | ≥6.7 | ≥6.7 | ≥6.7 | ≥6.7 |
| Clean | 6.0 | ≥6.5 | ≥6.5 | ≥6.5 | ≥6.4 | ≥6.4 | ≥6.4 | ≥6.4 |
| FBS | 0.3 | 1.1 | 4.6 | 4.8 | 0.3 | 0.7 | 3.9 | 4.7 |

### Efficacy testing on train parts

The efficacy of Zoono Z71 was also tested on a train tray table, arm rest and hand pole. Based on previous results, the efficacy of the antimicrobial coating was evaluated in the presence or absence of FBS. When results for the tray table and arm rest were analysed, there was a significant three-way interaction between the presence of the product, the presence of FBS and the surface to which the product was applied. The tray table comprised two different materials (high pressure laminate and stainless steel). In the absence of FBS, the average log_10_ reduction associated with the antimicrobial coating was significantly higher on stainless steel compared to high pressure laminate. The addition of FBS significantly reduced the efficacy of Zoono Z71 regardless of material (Supplementary table 6). When the coating was applied to high pressure laminate, and in the absence of interfering material, the concentration of ϕ6 was reduced by at least 4.8 log_10_ values within 120 min. The presence of FBS reduced the efficacy of the coating with log_10_ reductions not exceeding 1.2 over the same contact period. Similarly, when the coating was applied to a ‘clean’ and ‘dirty’ stainless steel surface, the concentration of ϕ6 after 120 min was reduced by ≥6.7 and <1.8 log_10_ values respectively (Table 3). Overall recovery of ϕ6 after 120 minutes was lower from the Terluran 22 armrest compared to other materials, but application of the antimicrobial coating did not lead to significant reductions of ϕ6 concentration on this material (Supplementary table 6). Regardless of the presence of FBS, <3log_10_ reductions were reported on all experiments (Table 3).

**Table 3:** Efficacy of Zoono Z71 when applied to train parts. Surfaces were coated with a layer of FBS prior to inoculation of ϕ6 and efficacy (calculated as mean log_10_ reduction (n=3)) was compared to when the product was applied to a ‘clean’ surface (i.e. when no interfering substance was present). Undetected virus observations were assumed to have a concentration of 20 PFU/replicate, the theoretical detection limit, for analysis. No data is available for cells shaded in grey.

| Days after coating was applied |  | Tray Table (high pressure laminate) |  | Tray Table (stainless steel) |  | Arm Rest (Terluran 22) |  | Hand pole (plastic coated) |  |
| --- | --- | --- | --- | --- | --- | --- | --- | --- | --- |
|  |  | Contact Time |  |  |  |  |  |  |  |
|  |  | 15 min | 120 min | 15 min | 120 min | 15 min | 120 min | 15 min | 120 min |
| ≤ 7 | Clean |  | ≥6.9 | 0.2 | ≥6.8 | 0.0 | 1.3 | 4.8 | 5.9 |
|  | FBS | 0.0 | 1.2 | 0.1 | 1.8 | 0.0 | 0.5 | 0.6 | 0.9 |
|  | Clean |  |  |  |  | 0.0 | -0.1 |  |  |
|  | FBS |  |  |  |  | 0.1 | 0.4 |  |  |
| >7 and ≤ 14 | Clean |  | 4.7 |  | ≥6.7 | -0.1 | 1.5 |  |  |
|  | FBS | 0.0 | 0.3 | 0.1 | 0.6 | -0.1 | 2.5 |  |  |
| >14 and ≤ 21 | Clean | 0.1 | 5.8 | 5.3 | ≥7.1 |  |  | 0.8 | ≥6.5 |
|  | FBS | 0.2 | 0.8 | 0.1 | 0.7 |  |  | 0.5 | 0.9 |
| >21 and ≤ 28 | Clean | 0.0 | 4.7 | 0.9 | ≥6.7 | 0.1 | 1.8 |  |  |
|  | FBS | 0.1 | 0.4 | -0.1 | 0.4 | 0.0 | 1.3 |  |  |

The results for the hand pole had to be analysed independently as a different inoculum size was used due to structural constraints. A significant interaction was found between presence of the antimicrobial coating and the presence of FBS. As with the tray table, coating the hand pole resulted in a significant reduction in ϕ6 (> 5 log_10_ reductions) after a contact time of 120 minutes but not in the presence of FBS (0.9 log_10_ reduction) (Table 3; Supplementary table 7).

The presence of FBS did not significantly affect the average log_10_ counts on any of the train surfaces when no antimicrobial coating was present (Supplementary tables 6 and 7).

The next study determined whether contaminating organic debris could be removed from a surface to restore the efficacy of the product. Coated and non-coated tray tables with and without FBS were wiped with a cloth wetted with sterile water prior to inoculation of ϕ6. A significant interaction was found between the application of the antimicrobial coating, the presence of FBS and whether the surface had been wiped (Supplementary table 8). As seen previously, when FBS was present, the efficacy of the product was depleted (<0.5 log_10_ reduction). Wiping the surfaces with a wetted cloth after FBS application did not restore the efficacy of the coating (Figure 3) with log_10_ reductions remaining <0.5 log_10_. When no interfering substance was present, wiping significantly reduced the efficacy of the coating (Supplementary table 8). In the absence of FBS, the efficacy of the coating was reduced after wiping the surface with the 10-wipe protocol; from 6.1 to 3.9 log_10_ reduction for high pressure laminate and from 5.6 to 0.9 log_10_ reduction for stainless steel. Increasing the number of wipes to 40 reduced the efficacy of the coating still further, from 5.3 to 2.2 log_10_ reduction for high pressure laminate (Figure 3).

**Figure 3:**
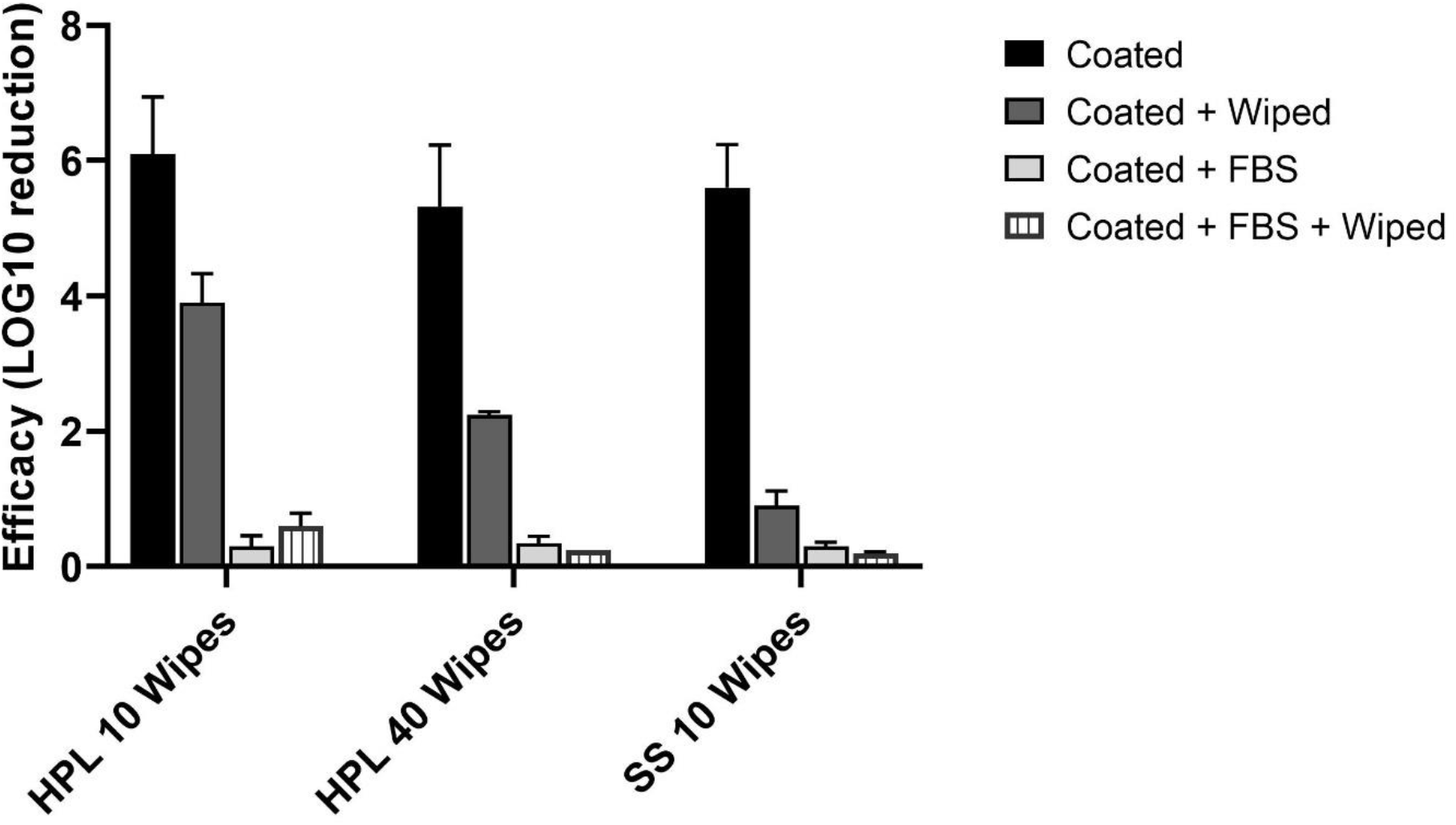
Efficacy of Zoono Z71 when applied to a tray table (HPL- high pressure laminate; SS-stainless steel), in the presence or absence of FBS, after wiping with a wetted cloth in comparison with non-wiped controls. Two wiping protocols differing in the number of wipes were applied. Efficacy was calculated as the average log_10_ reduction (n=2 with 3 replicates each) from ϕ6 pfu/replicate immediately after inoculation (time=0) minus ϕ6 pfu/replicate after 120 minutes. Errors bars represent standard deviation. Undetected virus observations were assumed to have a concentration of 20 PFU/replicate, the theoretical detection limit, for analysis.

## Discussion

Environmental surfaces can become contaminated by a range of pathogenic microorganisms and can contribute to the spread of infectious diseases, including respiratory viruses such as SARS-CoV-2. Strategies to reduce and contain shedding, such as hand hygiene and use of face coverings, and to reduce the microbial burden on surfaces, such as cleaning and disinfection, can prevent transmission through this route (17, 18). Antimicrobial coatings are an attractive strategy to supplement cleaning and maintain low levels of surface contamination between cleaning episodes but evidence of efficacy to support their implementation is lacking.

Quaternary ammonium compound-based disinfectants are widely used in industrial, healthcare, and domestic settings (19) and have been shown to be effective against SARS-CoV-2 (20). They are cationic detergents that present with a wide variety of chemical structures suitable for different applications, including polymer-based coatings, and have been shown to inactivate bacteria, yeast and viruses (19). In this and previous studies, when assessed under laboratory conditions and in the absence of interfering substances, quaternary ammonium polymer-based coatings have often shown high virucidal efficacy against SARS-CoV-2 and its surrogates (ϕ6 and human coronavirus 229E) when applied to stainless steel, polystyrene, Acrylonitrile Butadiene Styrene (ABS) plastic and poly(methyl methacrylate) coupons (14, 21, 22, 23, 24). However, it is worth noting that low efficacy has also been reported for some QAC-based products, even those with the same or very similar active ingredients as those previously shown to be effective (14, 23), suggesting that different formulations of QAC-based antimicrobial coatings do not necessarily share the same efficacy profile.

In this study, we extended the testing to other materials frequently used in the transport industry and observed different levels of efficacy. While the product was effective when applied to a high pressure laminate tray table and a plastic coated hand pole, high variability between replicates was observed when it was applied to glass and no virucidal activity was reported for the Terluran 22 armrest. Whilst the manufacturer’s claim the coating to be effective on all materials, these results suggest that antimicrobial coatings should be tested on all surface materials where application is intended as efficacy might differ among them. Hardison et al., 2021 (23) also observed high variability between replicates for one of the products they tested and differences in efficacy between application on stainless steel and ABC plastic were reported for another product. Although it has been claimed that QAC immobilized on a surface inactivates bacteria and viruses on contact (25), this does not mean that the effect is immediate. In this study and in the absence of interfering substances, contact times of at least 15 minutes were often required to achieve >3log_10_ reductions.

Public transport is a busy multi-user environment where organic debris frequently deposits on surfaces. This scenario was simulated by incorporating a layer of BSA or FBS over the coating. BSA is a protein commonly used in disinfectant testing (20, 26) and is the major component of FBS, which has also been previously used as an organic challenge (23, 24, 27). FBS has many more components including, lipids and carbohydrates and it is therefore likely to be more representative of general organic debris present on surfaces within public transport. In this study, to simulate accumulation of organic debris on surfaces, the interfering substances were applied as a layer over surfaces that had previously been coated with the antimicrobial, as opposed to adding the organic load to the viral suspension. While BSA did not interfere with the product, when a layer of FBS was applied to the test coupons, virucidal activity was reduced. When FBS was applied to the coated train parts the virucidal activity of the product was eliminated. Different application methods (coupons were coated by pipetting, while train parts were coated by fogging) or differences in the materials might explain the discrepancy between the reduction of virucidal activity on coupons and elimination on train parts. In agreement with these results, FBS has previously been reported to diminish the virucidal activity of a Benzalkonium chloride-based liquid disinfectant against equine herpesvirus type 1 (27).

As the product is designed to be used alongside regular cleaning, we also investigated whether wiping to remove the FBS from the tray table could restore the efficacy of the coating on this surface. When FBS was present, wiping with a wetted cloth was not able to restore the efficacy of the product. When FBS was not present, wiping with a wetted cloth reduced the virucidal activity of the product, suggesting that the coating was being removed from the surface by the mechanical action of cleaning. Whether the FBS had permanently inactivated the antimicrobial coating or if the coating was more effectively removed from the surface in the presence of FBS is unknown. Previous studies have found similar issues with the durability of QAC-based antimicrobial coatings. Butot et al. (21) reported loss of antiviral activity of a QAC-based antimicrobial coating after only one round of cleaning with a microfibre cloth and a water-based detergent or disinfection with 70% ethanol. Calfee et al. (14) found that seven QAC-based formulations effective against ϕ6 on initial testing were completely ineffective following exposure to wet abrasion cycles in accordance to the US EPA interim guidance (15). Other laboratory studies evaluating QAC-based products have not attempted to assess the durability (22-24).

This study has several limitations. Firstly, although tests were conducted using train parts taken from a scrap train and FBS was used to simulate organic debris, experiments were conducted under laboratory conditions which do not fully represent a real-life setting. While every effort has been made to identify the materials of the train parts used in this study, they might not be completely accurate, and they are unlikely to be representative of all materials used in public transport. Moreover, testing was done using the bacteriophage ϕ6 as a surrogate for SARS-CoV-2. ϕ6 is an enveloped dsRNA phage of the *Cystoviridae* family which shares structural features with SARS-CoV-2: it is enveloped by a lipid membrane, has spike proteins, is of similar size (∼ 80–100 nm) (28, 29) and has been previously used as a surrogate for coronaviruses in studies evaluating environmental persistence and disinfection efficacy (29-32). However, the suitability of ϕ6 as a surrogate will depend on the specific experimental conditions (30) and therefore the results on this report might not accurately represent the behaviour of SARS-CoV-2. Any materials associated with the virus (cell debris, soil, fluids) are also known to impact disinfection efficacy (33). Here we used TSB as a matrix (instead of respiratory fluids/sweat) for the viral suspension and also applied FBS to the surface (instead of organic debris found on public transport) which might have also influenced the results. High viral titres were used in this study and it is unknown whether virucidal activity would differ at lower titres. Finally, the ϕ6 suspension was applied as a 5-10 µL droplet which is representative of large droplet contamination but may not accurately represent deposition of virus for all contamination scenarios (e.g. by touch, smaller droplets or aerosols). Previous work with bacteria reported reduced efficacy for copper alloys when the inoculum was applied as an aerosol under reduced relative humidity (40%) as opposed to a wet inoculum (34). It is therefore possible that efficacy of the product has also been overestimated for some contamination scenarios in this study.

In conclusion, this study highlights the importance of testing QAC-based antimicrobial coatings on a range of materials, including relevant interfering substances and assessing durability in order to decide whether their application will reduce surface contamination burden in a given setting. For this particular product, the results suggest that efficacy in public transport surfaces will likely be quickly diminished due to the accumulation of organic debris which is inevitable in a busy, multi-user environment. Given that wiping the dirty surfaces with a wetted cloth failed to restore efficacy it is unlikely that application of this product will provide a substantial benefit in this setting. More research is warranted to identify suitable infection prevention and control practices for public transport.

## Acknowledgements

This work was supported by the TRACK: Transport Risk Assessment for COVID Knowledge project - EPSRC, EP/V032658/1-. We would like to thank Jonathan Bridgewood and First Group for providing the train parts. The views expressed in this article are those of the authors and are not necessarily those of UKHSA or the Department of Health and Social Care.

